# Neural coding of visual objects rapidly reconfigures to reflect sub-trial shifts in attentional focus

**DOI:** 10.1101/2021.05.25.445712

**Authors:** Lydia Barnes, Erin Goddard, Alexandra Woolgar

## Abstract

Every day, we respond to the dynamic world around us by flexibly choosing actions to meet our goals. This constant problem solving, in familiar settings and in novel tasks, is a defining feature of human behaviour. Flexible neural populations are thought to support this process by adapting to prioritise task-relevant information, driving coding in specialised brain regions toward stimuli and actions that are important for our goal. Accordingly, human fMRI shows that activity patterns in frontoparietal cortex contain more information about visual features when they are task-relevant. However, if this preferential coding drives momentary focus, for example to solve each part of a task, it must reconfigure more quickly than we can observe with fMRI. Here we used MVPA with MEG to test for rapid reconfiguration of stimulus information when a new feature becomes relevant within a trial. Participants saw two displays on each trial. They attended to the shape of a first target then the colour of a second, or vice versa, and reported the attended features at a choice display. We found evidence of preferential coding for the relevant features in both trial phases, even as participants shifted attention mid-trial, commensurate with fast sub-trial reconfiguration. However, we only found this pattern of results when the task was difficult, and the stimulus displays contained multiple objects, and not in a simpler task with the same structure. The data suggest that adaptive coding in humans can operate on a fast, sub-trial timescale, suitable for supporting periods of momentary focus when complex tasks are broken down into simpler ones, but may not always do so.

---

Human cognition is remarkably flexible. We can fluidly direct our focus towards what we need for our current goal, seamlessly adapt to changes in our environment, and generalise from what we know to solve new problems. Several lines of research suggest that this flexibility emerges from activity in frontoparietal cortex. Cognitively challenging tasks elicit robust activity in the ‘multiple demand’ (MD) system—a distributed network of frontal and parietal cortex recruited by a wide range of tasks (Assem et al., 2020; Duncan, 2010; Fedorenko et al., 2013). Damage to this system linearly predicts fluid intelligence scores (Woolgar et al., 2010, 2018), which in turn powerfully predict how well we are able to acquire new skills.

The characteristic adaptability of MD regions means that they are ideally suited to supporting flexible cognition. Patterns of activity in the MD system, measured with fMRI, adapt to code information that is relevant for the current task. MD patterns can encode many different aspects of a task (for example, visual: Jackson et al., 2016; vibrotactile: Woolgar & Zopf, 2017; for a review see Woolgar et al 2016), commensurate with a high degree of mixed selectivity in these regions (Fusi et al., 2016; Rigotti et al., 2013). Moreover, MD coding for task-relevant stimuli is enhanced when stimuli are more difficult to discriminate (Woolgar, Hampshire, et al., 2011; Woolgar, Williams, et al., 2015) and changes to prioritise information that is at the focus of attention (Jackson & Woolgar, 2018; Woolgar, Williams, et al., 2015). Activity in at least one MD region appears to be causal for facilitating task-relevant information processing (Jackson et al., 2021). This, in turn, may provide a source of bias to more specialised brain regions, for example through task-dependent connectivity (Cole et al., 2013; see for example Baldauf & Desimone, 2014). Consequently, adaptive coding has been proposed as a central component of goal-directed attention, biasing sensory and motor brain regions to perceive and respond to information that is relevant to our current task.

A key outstanding question concerns the temporal scale of this process. Here, we explore the ‘attentional episodes’ account of flexible behaviour (Duncan, 2013) which predicts a fast temporal scale. This account draws on studies of human and artificial intelligence to propose that flexible behaviour rests on our ability to break a complex task down into a series of simpler parts, and to focus, moment-to-moment, on the information needed for each part (Duncan, 2013; Duncan et al., 2012, 2017). Indeed, there is some evidence that this ability may underpin performance on novel problem solving tasks. For example, explicitly breaking a complex task into simple parts removes the performance gap between people with high and low fluid intelligence scores (Duncan et al., 2017; see also O’Brien et al., 2020). In this matrix reasoning study, participants viewed a 2×2 grid with three of the four squares filled with an image. They had to abstract relationships between the images to fill in the remaining square. Images consisted of multiple features. In the second half of the experiment, each feature was presented separately. These segmented problems were trivial to solve, regardless of whether participants struggled or performed well on the difficult, unsegmented problems. This led the authors to propose that participants who were able to solve the unsegmented problems were better able to mentally break them down into their relevant parts. Adaptive coding could be a key component of this segmentation by driving momentary focus toward subsets of the available information in turn.

From these studies, it seems intuitive that flexible cognition involves identifying simple problems that we can solve, and addressing them in an ordered sequence. However, we do not have clear insight into how adaptive codes reconfigure to prioritise relevant information throughout a task. The bulk of research on adaptive coding in humans uses fMRI. While these studies show trial-to-trial shifts in what information can be discriminated from activity patterns (for example, Woolgar et al., 2011, 2015), the coarse temporal resolution of fMRI does not support precise, sub-second measurement of changes in task information.

Time-resolved neuroimaging methods, by contrast, offer promising evidence for rapid changes in task representation. Non-human primate studies show that the same frontal neurons can encode object identity, then location, within a single trial, as monkeys attended to what, then where, an object was (Rao et al., 1997). Although these data are taken from highly trained monkeys and could rely on a learned response rather than instantaneous shifts in a flexible brain system, they demonstrate that the neural population can systematically change its activity pattern in synchrony with the task. More recent work by Spaak et al. (2017) demonstrates that, even when the same information is encoded across phases of a task, neurons in primate lateral prefrontal cortex dynamically update what they encode. This dynamic reallocation of selectivity within a trial makes plausible rapid shifts in the information that these adaptive brain regions represent. In humans, stronger coding for visual features when they are task-relevant compared to task-irrelevant, emerges in MEG data as early as 100 ms from stimulus onset (Battistoni et al., 2018; Goddard et al., 2019; Moerel et al., 2021; Wen et al., 2019), with sustained coding of the relevant feature emerging around 200-400 ms in the MEG/EEG signal (Goddard et al., 2019; Grootswagers et al., 2021; Moerel et al., 2021; Yip et al., 2021). This provides preliminary evidence that population codes for task-relevant features develop rapidly.

However, this previous time-resolved human neuroimaging work did not require participants to shift their attention within trials, so we do not know how rapidly information codes update to redirect attention in each part of a task. This is important to study, however, if it is indeed a key component of how we solve complex tasks. Here, we test the dynamic adaptation of task representations as what is relevant changes within single trials. We use MEG to track shifts in adaptive coding with sub-second precision across fragments of two rapidly changing tasks. Considering the strong association between task difficulty and the brain regions implicated in adaptive coding (Crittenden & Duncan, 2014; Fedorenko et al., 2013), we test this at two levels of attentional demand. In Experiment 1, we use simple stimuli to track preferential coding of relevant information under low attentional demands. In Experiment 2, we use a complex stimulus space, abstracted decisions, and the presence of distractors to track preferential coding of relevant information under high attentional demands. Across both experiments, we ask whether neural codes for relevant stimulus information rapidly reconfigure when what is relevant changes mid-trial.

## Methods

### Participants

Participants were selected to (a) have normal or corrected-to-normal vision (must have normal colour vision); (b) be right handed; (c) have no exposure to fMRI in the previous week; (d) have no non-removable metal objects; and (e) have no history of neurological damage or current psychoactive medication. Prospective participants were informed of the study’s selection criteria, aims, and procedure, through a research participation site.

For Experiment 1, 20 participants (17 female, 3 male, mean age 25±6 years) were recruited from the paid participant pool at Macquarie University (Sydney). They gave written informed consent before participating, and were paid AUD$30 for their time. Ethical approval was obtained from the Human Research Ethics Committee at Macquarie University (5201300602).

For Experiment 2, 20 participants (16 female, 4 male, mean age 31±12 years) were recruited from the volunteer panel at the MRC Cognition and Brain Sciences Unit (Cambridge). They gave written informed consent prior to each testing session, and were paid GBP£40 for their time. Participants were additionally asked to only volunteer if they had existing structural MRI scans on the panel database. Two participants took part prior to completing a structural scan; one obtained a scan through another study conducted at the MRC Cognition and Brain Sciences Unit, while the other returned for a separate MRI session as part of this study. This participant gave written informed consent before completing the structural scan and was paid an additional GBP£20 for this component of their time. Ethical approval was obtained from the Psychology Research Ethics Committee at the University of Cambridge (PRE.2018.101).

### Stimuli

Stimuli were created in MATLAB and presented with Psychtoolbox (Brainard, 1997; Kleiner et al., 2007). In Experiment 1, they were displayed with an InFocus IN5108 LCD back projection monitor (InFocus, Portland, Oregon, USA) at a viewing distance of 113 cm. In Experiment 2, they were displayed with a Panasonic PT-D7700 projector at a viewing distance of 150 cm.

Experiment 1 stimuli consistent of four novel objects (Op de Beeck et al., 2006; see Figure 1) that were either ‘cubie’ or ‘smoothie’ shaped, and green or red (RGB 0-194-155 and 224-0-98). Colours were chosen for high chromatic variation and strong contrast against the dark grey background (RGB 30-30-30).

**Figure 1.**
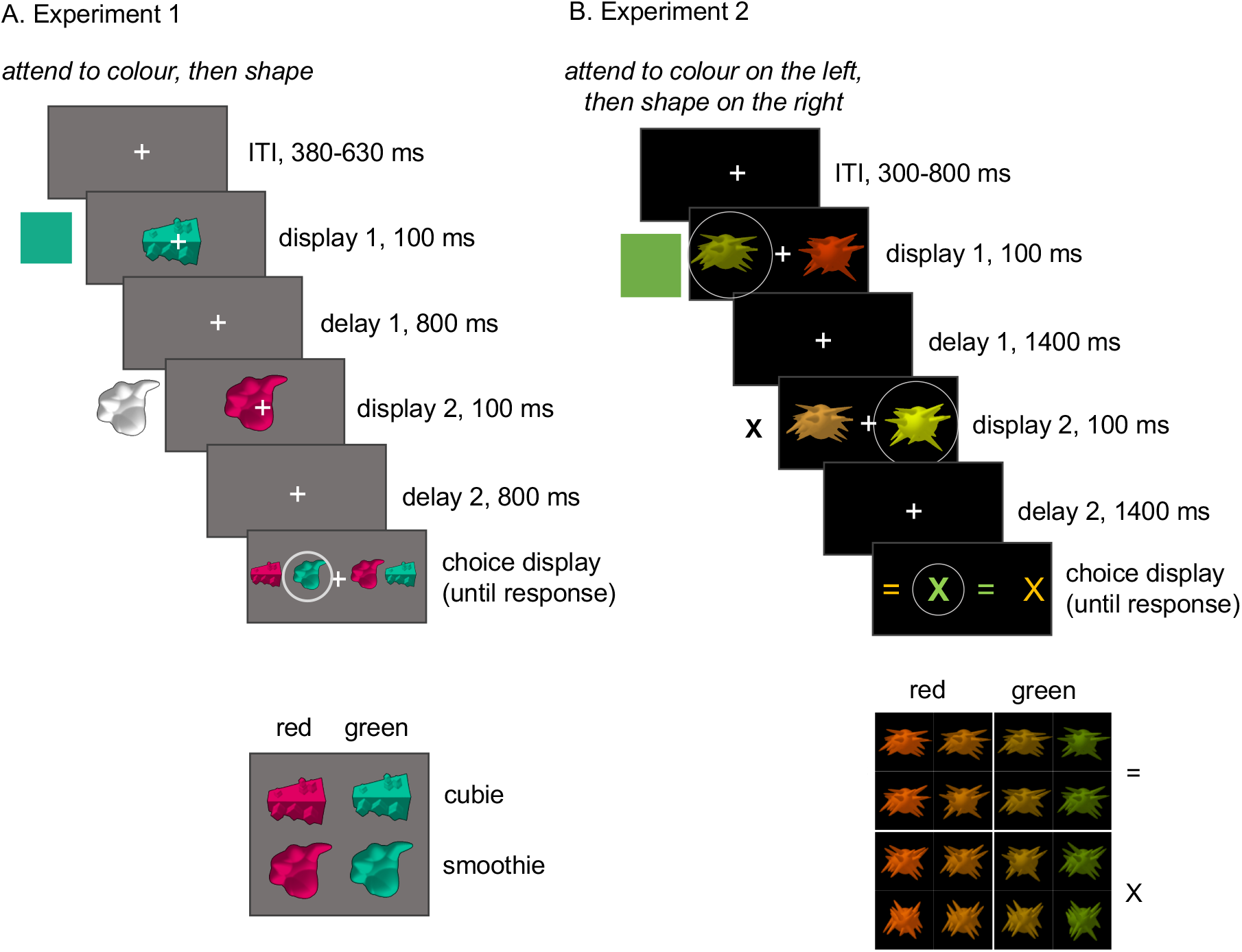
Stimuli and example trials for Experiment 1 and 2. Relevant information for each epoch is shown beside the display. Panel A shows an example trial for Experiment 1, with a single object on each display. In this trial, the relevant features are “green” (Target 1) and “smoothie” (Target 2), resulting in a “green smoothie” response on the choice display. Stimuli could be red, green, “cubie”, or “smoothie”. Panel B shows an example trial for Experiment 2, in which the participant was cued to attend to colour on the left, then shape on the right. The relevant features were thus green and “X”, leading to a response of “green X” on the choice display. Stimuli varied in four steps from red to green, and from X to =, but were assigned to binary red / green, X / = categories. Circles represent the focus of attention and correct choice and were not shown to participants.

Experiment 2 stimuli consisted of 16 novel “spiky” objects, adapted from the Op de Beeck et al (2006) ‘spiky’ stimuli, selected at four points on a spectrum of red to green, and upright to flat (Goddard et al., 2019). Colour values were numerically equally spaced in u’v’ colour space between [0.35,0.53] and [0.16,0.56]. Shapes were also equally spaced to create four steps in orientation from upright to flat. Each step included 100 shape exemplars, with different spikes indicating the orientation, to discourage participants from judging orientation based on a single spike.

### Task

Experiment 1 used simple displays and stimuli, optimised for strong visual signals. Each block began with a written cue instructing participants to attend to the colour of the first object, and the shape of the second object, or vice versa. On each trial, participants viewed two brief displays (100 ms), each followed by a delay (800-1500 ms; see Figure 1). Finally, they were prompted to select an object from a choice display that comprised the combination of the remembered features. All four objects appeared on the choice display, and participants selected the object that matched the colour and shape they had extracted from the preceding displays. For example, under the rule “attend shape, then colour”, if the first object was a ‘cubie’ and the second object was ‘red’, the target on the choice display was a red cubie. Participants indicated their choice by pressing one of four buttons on a bimanual fibre optic response pad operated with the four fingers of the right hand. The mapping from object location to response button was intuitive (far left button for far left object, etc) and consistent across trials; however, the arrangement of the four objects on the choice display varied to prevent participants preparing a motor response until the display screen was shown. Stimulus arrangements were presented in pseudo-random order and balanced within each rule such that all stimuli on the second display were equally preceded by each stimulus on the first display, and the correct choice pertained equally to all motor responses. If a participant made three consecutive incorrect or slow responses (>3 s), the task was paused and the cue was presented again until the participant verbally confirmed that they understood the rule for that block. Average accuracy and response times were displayed at the end of each block.

Experiment 2 followed the structure of Experiment 1, but used simultaneously presented objects and subtle stimulus discriminations, optimised for high attentional load. For this experiment, each display contained two objects. Participants were cued to both a location and feature, for example, “attend to shape on the right, then colour on the left”. Relevant location and feature always changed from display 1 to display 2, creating four possible rules. Participants judged the colour and shape *category* of the target objects’ features. The choice display contained the symbols X and =, presented in the average of the two ‘red’ colours and the average of the two ‘green’ colours, to represent the four possible answers. These symbols were chosen to encourage participants to make category-level decisions about the objects.

### Procedure

#### Experiment 1

Each participant first completed four blocks of 10 practice trials outside the shielded room. These were identical to test trials except that (a) participants received feedback of ‘Correct’, ‘Incorrect’, or a red screen signifying a slow response (>3 s), on every trial, (b) display durations in the first two practice blocks were slowed from 100 ms to 500 ms to ease participants into the task, and (c) response key codes were marked on the choice display to train participants in the location-response mapping. Once in the MEG scanner, participants completed eight blocks of 96 trials each, with feedback at the end of each block. Each block lasted approximately seven minutes. Blocks alternated between the two rules, ‘attend shape, then colour’ and ‘attend colour, then shape’, with the order counterbalanced across participants.

#### Experiment 2

Participants learned the stimulus categories (red vs green, upright vs flat) and the task in a separate training session. Training could be on the day of or the day before the scanner session. Training consisted of two blocks of 50 category learning trials, in which they saw a single object for 100 ms and pressed a button to indicate its shape or colour category. They then began training on the core task. Within-trial delay periods began at 4 s and reduced to 1.5 s in three steps (3 s, 2 s, 1.5 s). Participants completed a minimum of 10 trials at each of the four speeds for each of the four rules (that is, at least 40 trials per rule).

After 10 trials were completed, the speed increased when the participant got 8 trials correct in any 10 consecutive trials. Feedback was given on each trial by a brighter fixation cross for correct responses and a blue fixation cross for incorrect responses, shown for the first 100 ms of the post-trial interval. This procedure trained each participant to the same criterion without penalising them for errors early in the block.

Once in the MEG, participants completed four blocks, each corresponding to a single rule and comprising 258 trials, lasting approximately 20 minutes. Rule order was balanced across participants.

### MEG data acquisition

#### Experiment 1

We acquired MEG data in the Macquarie University KIT-MEG lab using a whole-head horizontal dewar with 160 coaxial-type first-order gradiometers with a 50 mm baseline (Model PQ1160R-N2; KIT, Kanazawa, Japan; Kado et al., 1999; Uehara et al., 2003) in a magnetically shielded room (Fujihara Co. Ltd., Tokyo, Japan). First, the tester fit the participant with a cap containing five head position indicator coils. The location of the nasion, left and right pre-auricular, and each of the head position indicators were digitised with a Polhemus Fastrak digitiser (Polhemus, VT, USA). This information was copied to the data acquisition computer to track head position during data collection. Participants lay supine during the scan, and were positioned with the top of the head just touching the top of the MEG helmet. Any change in head position relative to the start of the session was checked and recorded after four blocks. MEG data were recorded at 1000 Hz.

#### Experiment 2

We acquired MEG data with the MRC Cognition and Brain Sciences’ Elekta-Neuromag 306-sensor Vectorview system with active shielding. Ground and reference EEG electrodes were placed on the cheek and nose. Bipolar electrodes for eye movements were placed at the outer canthi, above and below the left eye. Heartbeat electrodes were on the left abdomen and right shoulder. Scalp EEG were also applied for a separate project. Head position indicators were placed on top of the EEG cap. Both head shape and the location of the head position indicators were digitised with a Polhemus Fastrak digitiser. Head position was recorded continuously through the scan and viewed after each block to ensure that the top of the participant’s head stayed within 6 cm of the top of the helmet in the dewar (mean movement across task 3.94 mm, range 0.5:15 mm). Because targets in this experiment could appear to either side of fixation, we also recorded eye-movements with an EyeLink 1000 eye tracker, which we calibrated before each block. If we observed more information about the stimulus at the relevant location, eye-tracking data would allow us to regress out the contribution of gaze. However, we did not observe a main effect of spatial attention, and so did not include the eye-tracking data in this analysis.

### Analyses

#### MEG processing

Due to active shielding and artefacts from continuous head position indicators, data from Experiment 2 were first processed with Neuromag’s proprietary filtering software (*Maxfilter*, 2010). We applied temporal signal space separation to remove environmental artefacts, used continuous head position information to correct for head movement within each block, and reoriented each block to the subjects’ initial head position.

All other processing was the same across experiments. We used a minimal preprocessing pipeline to minimise the chance of removing meaningful data. This was especially appropriate in our case, as our planned multivariate analyses are typically robust to noise (Grootswagers et al., 2016). MEG data were imported into MATLAB v2018b using Fieldtrip (Oostenveld et al., 2011), and bandpass filtered (0.01-200 Hz). Trials were epoched from a 100 ms pre-stimulus baseline to the maximum possible trial duration (Exp 1: 4800 ms, Exp 2: 5000 ms). All sensors were included in the initial analysis.

#### MEG decoding

We used multivariate pattern analysis to trace the information about rule, colour, and shape in each task phase. We then compared the information about colour when it was relevant and irrelevant, repeating the comparison for shape. Following previous studies, we expected that rule information, which was known before each trial, would be present throughout the trial and increase briefly after visual displays (Goddard et al., 2019; Hebart et al., 2018). We predicted that preferential coding would be reflected in improved decoding of visual features when they were relevant, compared to irrelevant (Battistoni et al., 2018; Goddard et al., 2019; Grootswagers et al., 2021; Hebart et al., 2018; Moerel et al., 2021; Wen et al., 2019; Yip et al., 2021). Increased colour information when colour was relevant would indicate that information was flexibly coded according to task demands. Our critical comparison, then, was how this happened for the two task phases. If information about the relevant feature was prioritised in both task epochs, this would indicate that preferential coding can reconfigure in line with sub-second shifts in what is relevant to the task.

We first trained a linear classifier (linear discriminant analysis, LDA; see Grootswagers et al., 2016) on labelled data from two feature rules—“attend colour, then shape” and “attend shape, then colour”—using all but one trial from each category. We then tested whether the weights that the classifier had learned to discriminate the training data generalised to the remaining unobserved trials. We repeated the process, leaving out a different pair of trials each time, until all trials had acted as the test data. We then averaged the classification accuracy across all test sets. For Experiment 2, we collapsed the feature rule analysis over locations to mirror Experiment 1. We also decoded the location rule (i.e., “attend left, then right” and “attend right, then left”), which we report in the Supplementary Materials (Figure S1) for completeness.

For colour and shape classification, we trained a linear classifier on labelled data from two categories—for example, “red” and “green”—using all but one trial from each category, for each feature rule separately. For Experiment 2, we decoded pairs of shape or colour, at a fixed location, for each feature and location rule. For example, we took trials under the rule “attend colour on the left, then shape on the right”. For items on the left on the first display, we decoded strong red vs yellow red, yellow red vs yellow green, and so on for all six pairs of colour. We then averaged classifier accuracy across the six pairs into a single measure of colour information coding in the left hemifield under this rule. We repeated this for each rule to obtain four traces of left hemifield colour information coding, representing colour information when that location and feature were relevant or irrelevant. We conducted the same pairwise decoding and averaging for colour in the right hemifield. Conducting the analyses for each hemifield separately minimised the requirement for the classifier to generalise patterns over space. Finally, we averaged the four traces of left hemifield colour information coding with the corresponding right hemifield traces to produce a single trace for each attention condition: “attended location, attended feature” (the task-relevant trace), “attended location, unattended feature”, “unattended feature, attended location”, and “unattended location, unattended feature”. The two traces for colour (or shape) information at the attended location parallel the two traces for each target in Experiment 1 and form the central part of our analysis.

#### Statistical tests

We tested whether decoding accuracy scores were above chance using a null distribution generated from the data. To generate this, we permuted the predicted class labels so that they were randomly assigned over trials (Bae & Luck, 2019). We calculated decoding accuracy as above and repeated the process 10,000 times to produce a decoding distribution for each participant and each comparison. We then sampled 10,000 times across participants’ null distributions to form a group-level null distribution. At each timepoint, we calculated t-scores for classification accuracy relative to the null distribution (Stelzer et al., 2013). We used a threshold-free cluster statistic (threshold step 0.1; Smith & Nichols, 2009) to flexibly set a cluster-forming threshold to identify peaks in the t-score timecourse that were more strong and/or sustained than expected from the null distribution (p<.05). This maximises sensitivity to peaks that are most likely to reflect meaningful change while down-weighting peaks that are small or transient (Smith & Nichols, 2009). We then used this threshold to correct for multiple comparisons at the cluster level across the whole trial. Decoding onset was the onset of the first cluster for which decoding accuracy was reliably above chance.

For between-condition comparisons, we contrasted the decoding trace for the target when it was the relevant or irrelevant feature using a two-sided t-test, implemented in CoSMoMVPA (Oosterhof et al., 2016) with threshold-free cluster enhancement and a threshold step of 0.1(p<.05; Smith & Nichols, 2009).

For Experiment 2, we also conducted secondary analyses to assess the combined effects of spatial- and feature-selective, as reported in Goddard et al. (2019). We conducted 2×2 ANOVAs to test, for each time bin, whether stimulus colour and shape information coding was boosted (1) at the relevant compared to irrelevant location, (2) when that stimulus feature was relevant for the task compared to when it was irrelevant, and (3) when both feature and location were relevant compared to all other attention conditions. We quantified these as main effects of spatial and feature-selective attention, and as a planned comparison between the coding of the reported feature at the attended location and the coding of that feature at that location in the other 3 attention conditions (following our prediction from Goddard et al., 2019). For example, we contrasted decoding for colour on the left when people were attending to colour on the left, with decoding for colour on the left when attending to shape on the left, colour on the right, and shape on the right. We present the results of these secondary analyses in the Supplementary Materials (Figure S2).

Lastly, in Experiment 2 we asked whether attentional effects had similar temporal profiles in epoch 1 and epoch 2 of the trial. We epoched the stimulus decoding traces for the target, separately around the first and second stimulus displays (0:1500 ms), using the same pre-trial baseline (−100:0 ms) for all traces. This created four overlaid traces, a relevant and an irrelevant feature trace for Epoch 1 and Epoch 2. We conducted a 2×2 ANOVA with main effects of relevance and epoch. An interaction term tested our hypothesis that preferential coding of relevant information emerges earlier, or is more substantial, in one epoch compared to the other.

## Results

### Behavioural performance

In Experiment 1, median accuracy was 93.3% (std 7.5%), with median reaction time 829.2 ms (std 210.7 ms). In Experiment 2, median accuracy was 75.9% (std 10.9%), with median reaction time 665.2 ms (std 92.1 ms). In both tasks chance accuracy would have been 25%.

### Rule information coding

We trained a classifier to discriminate between feature attention rules (“attend shape, then colour” from “attend colour, then shape”) to extract a time-course of rule information coding (Figure 2). Since the rule was cued at the start of the block, we expected that participants might prepare their task set in advance of the stimulus display. We anticipated that rule information would be more decodable after each display, when the rule could be applied to extract relevant information (as in Goddard et al., 2019). Based on this we predicted that we would be able to decode the rule from the pattern of MEG activity throughout the trial, with peaks during the early stimulus-induced responses (~100-300 ms after each stimulus onset). Indeed, rule information coding emerged early in both experiments, increasing after each stimulus onset, and remaining above chance throughout the trial. Rule information coding gradually ramped up after each display in Experiment 1, whereas in Experiment 2 rule information coding was elevated throughout the trial and peaked steeply after each display.

**Figure 2.**
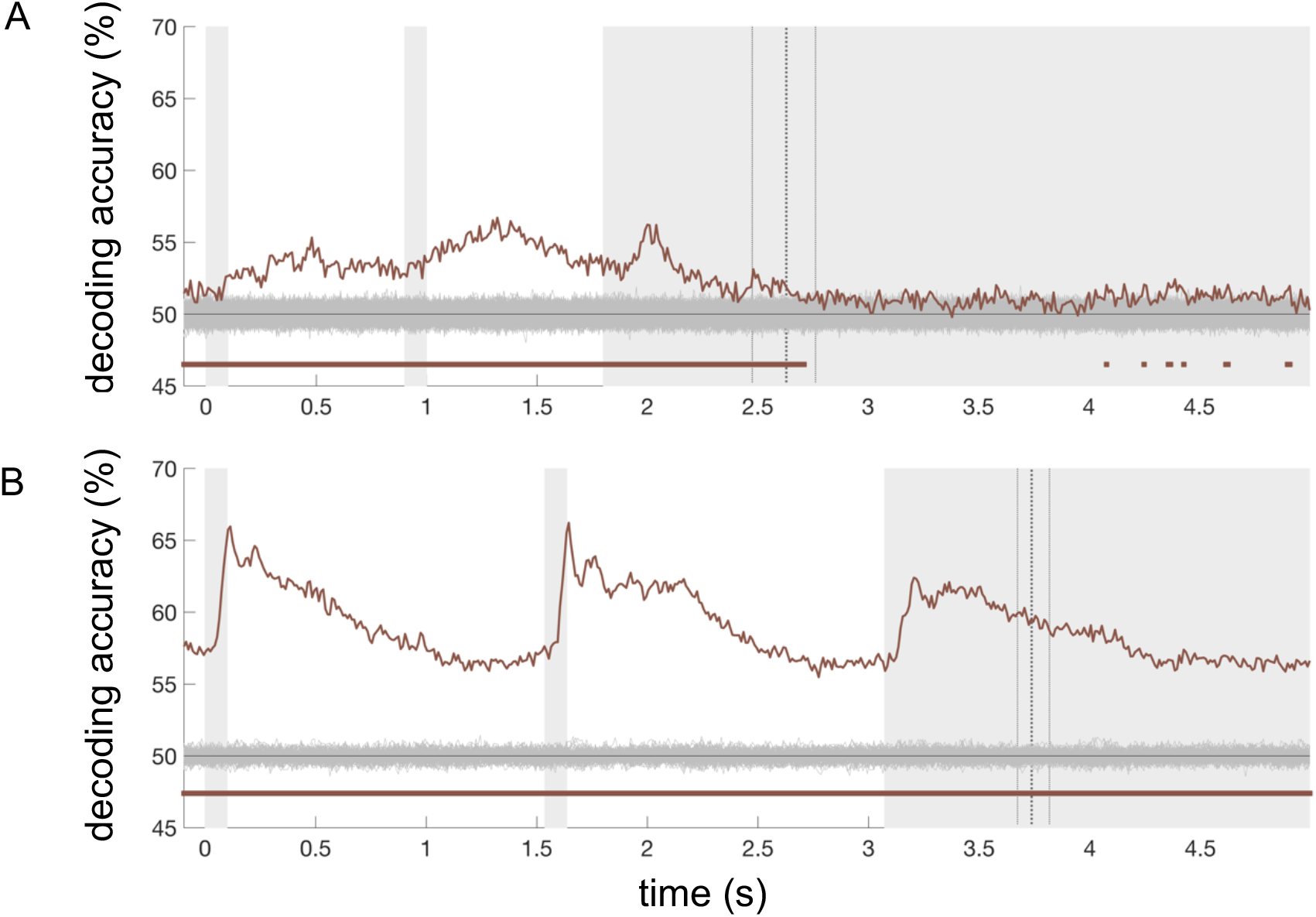
Feature rule decoding (“attend colour then shape” vs “attend shape then colour”) for Experiment 1 (A) and Experiment 2 (B). Vertical grey patches mark the stimulus displays and the maximum possible duration of the choice display. Vertical dotted lines mark the median response time with one quartile on either side. Permutation-based null data are shown in grey around chance (50%). Timepoints at which decoding was reliably different to the null based on threshold-free cluster correction are marked below the trace in brown.

### Preferential coding of visual features

Next, we examined the time-course with which we could decode stimulus colour and shape from the pattern of MEG activity (Figures 3 & 4). We quantified this separately when a feature was relevant or irrelevant for the participant’s task so that we could examine the effect of attention on coding of this information. We predicted that both relevant and irrelevant stimulus features would be decodable from the sensor data, but that each feature would be more readily decoded when it was relevant compared to when it was irrelevant (Goddard et al., 2019; Hebart et al., 2018; Moerel et al., 2021). In Experiment 1, robust decoding of stimulus information emerged rapidly after the onset of each display, remaining through the initial part of the delay phase for each epoch. Contrary to our prediction, however, in Experiment 1 there was no reliable evidence of preferential coding of the currently relevant information, in either task epoch, for colour or shape information (Figure 3).

**Figure 3.**
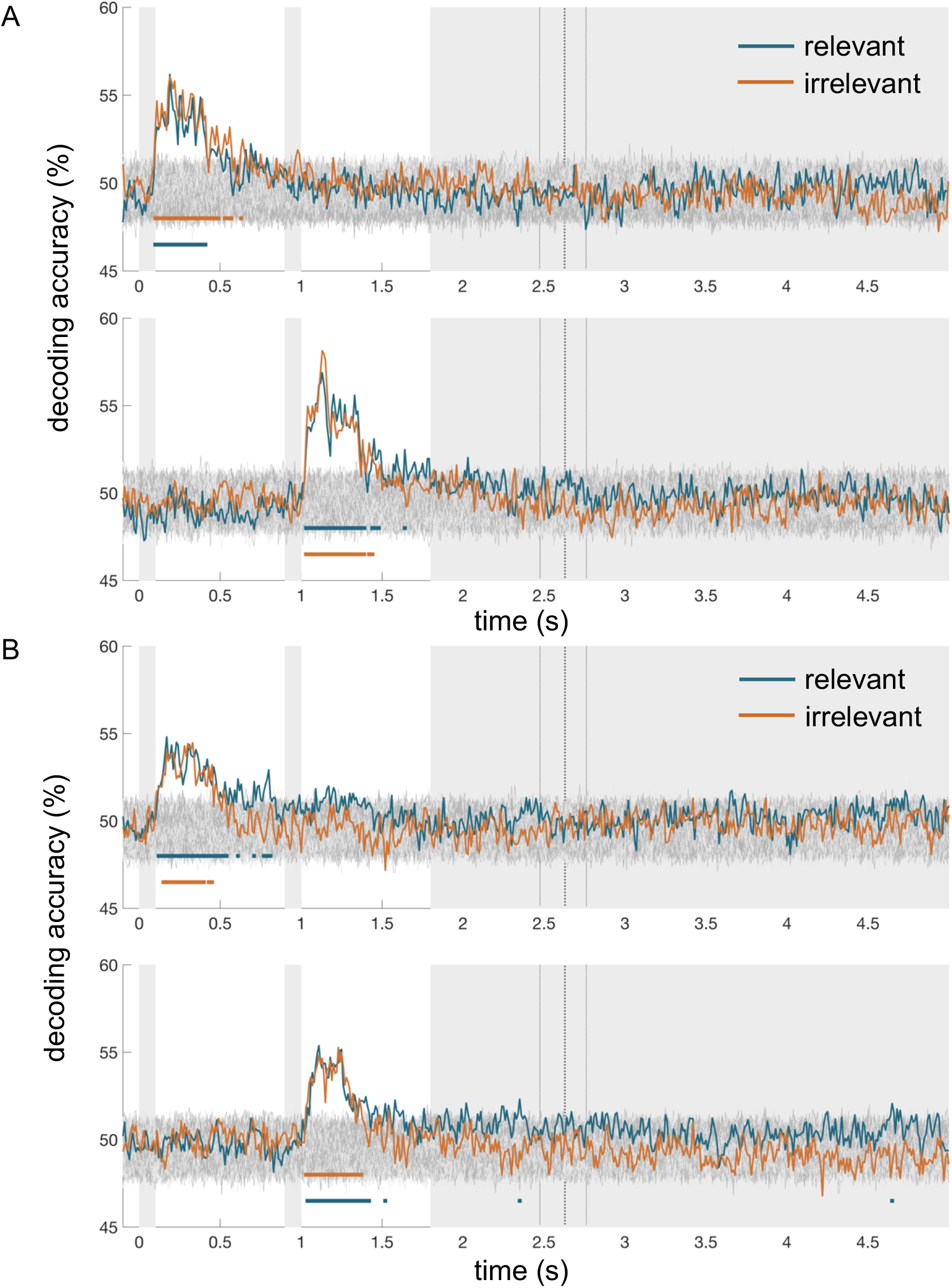
Colour (A) and shape (B) decoding for Experiment 1. A and B show decoding traces for the first and second targets in the upper and lower panels. Decoding accuracies are shown for each feature when it was relevant (blue) or irrelevant (orange) for the task. Grey bars mark the stimulus and response display durations. Vertical lines show the median response time, ± one quartile. Times at which decoding was greater than chance, p<0.05 using a cluster-based correction for multiple comparisons, are marked below each trace in the corresponding colour. Relevant information coding did not reliably exceed coding for the irrelevant feature at any timepoint.

Experiment 2 stimulus decoding was similarly rapid. Although less pronounced (potentially due to the busier displays and more subtle colour and shape differences) initial stimulus decoding peaks followed a similar timecourse to Experiment 1. For coding of colour, there was an initial stimulus-driven response peaking at 100 ms, which was similar when that information was relevant or irrelevant, and which occurred for both epochs, though these peaks did not reach statistical significance. For shape, the pattern was broadly similar and statistically significant, with an initial stimulus-driven response at 100 ms from each display onset. Critically, in contrast to Experiment 1, in Experiment 2 we now saw evidence of additional, sustained, preferential coding of relevant information. Whereas decoding for the target’s colour remained close to chance when that feature was irrelevant, coding for the same information when it was relevant was higher and sustained (Figure 4). Coding of relevant colour information was reliably different to chance and to the irrelevant feature trace from approximately 500 ms after stimulus presentation and was sustained into the subsequent trial epoch. We observed the same pattern for shape decoding, with a sustained response only for the relevant information in both epochs, which was significantly greater than the other attention conditions from approximately 500 ms after the display. Secondary analyses of all four stimulus traces (relevant and irrelevant features of target and distractor objects) showed a brief stimulus-driven peak followed by sustained preferential coding of the relevant target feature compared to the average of all other features, with inconsistent main effects of spatial or feature-selective attention (Figure S2).

**Figure 4.**
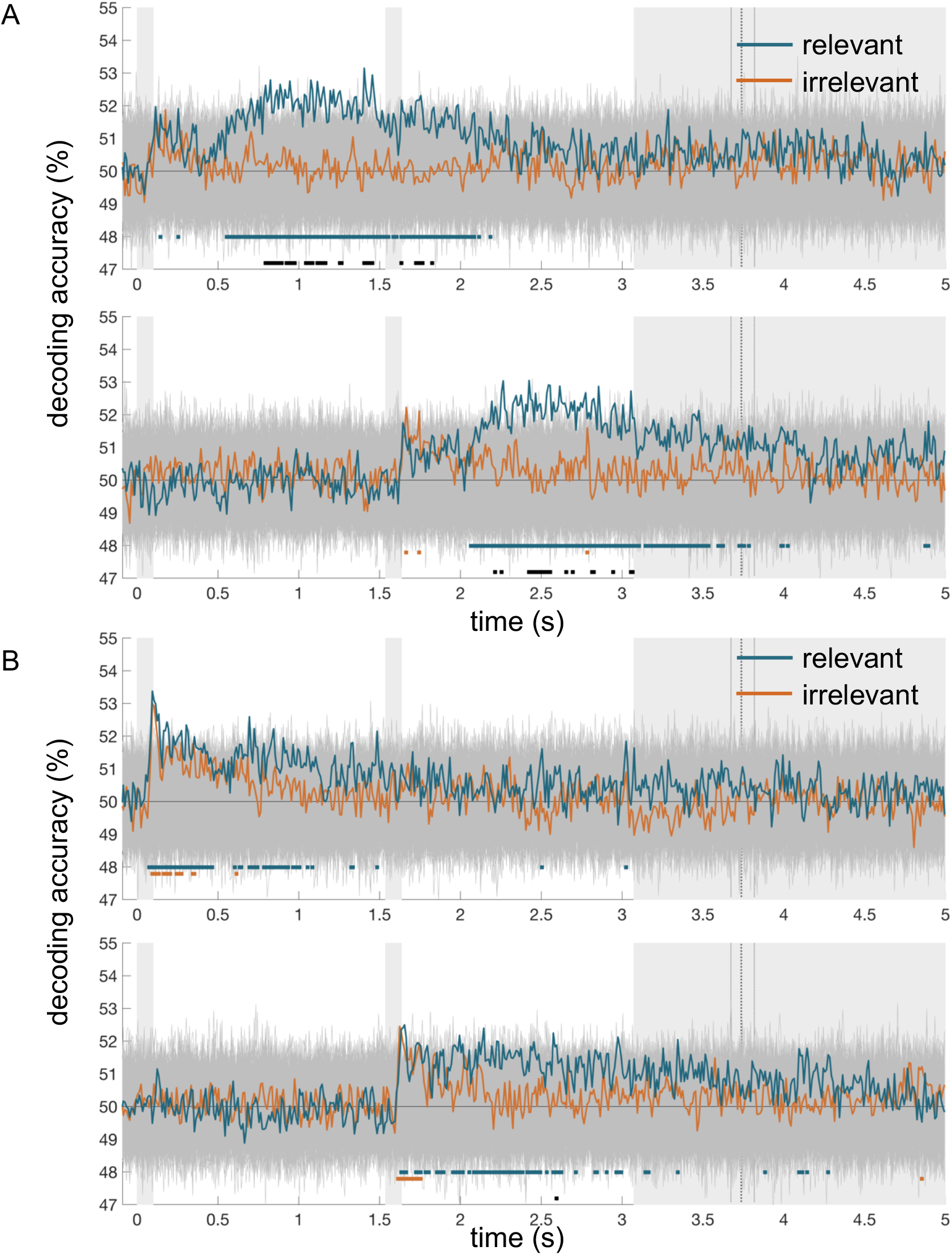
Colour (A) and shape (B) decoding for Experiment 2. A and B show decoding traces for the first and second targets in the upper and lower panels. Decoding accuracies are shown for each feature when it was relevant (blue) or irrelevant (orange) for the task. Grey bars mark the stimulus and response display durations. Vertical lines show the median response time, ± one quartile. Times at which decoding was greater than chance, p< 0.05, using a cluster-based correction for multiple comparisons, are marked below each trace in the corresponding colour. Times at which relevant information coding was reliably above coding for the irrelevant target feature (threshold-free cluster correction, p<.05) are marked in black.

### Rapid coding of features across epochs

To compare the dynamics of attentional prioritisation in the two epochs, we took the decoding traces for the target in each epoch of Experiment 2 and aligned them in time. We anticipated that the effect of attention (enhancement of relevant information) might develop later in Epoch 2, which reflected a sub-trial shift of attention when participants had less time to prepare what they would attend to. However, preferential coding for relevant information in Epoch 2 was comparable to Epoch 1 (Figure 5). We did not observe a main effect of epoch, or an interaction between epoch and relevance. This does not rule out the possibility that shifting attention mid-trial incurs some delay in preferential coding, for example in other situations. However, it demonstrates that humans can rapidly reconfigure their neural codes to prioritise coding of a new stimulus dimension mid-trial, even while holding the previously attended stimulus information in mind. Reinforcing non-human primate work, this highlights that our capacity to adaptively code task-relevant information is flexible over time as well as content.

**Figure 5.**
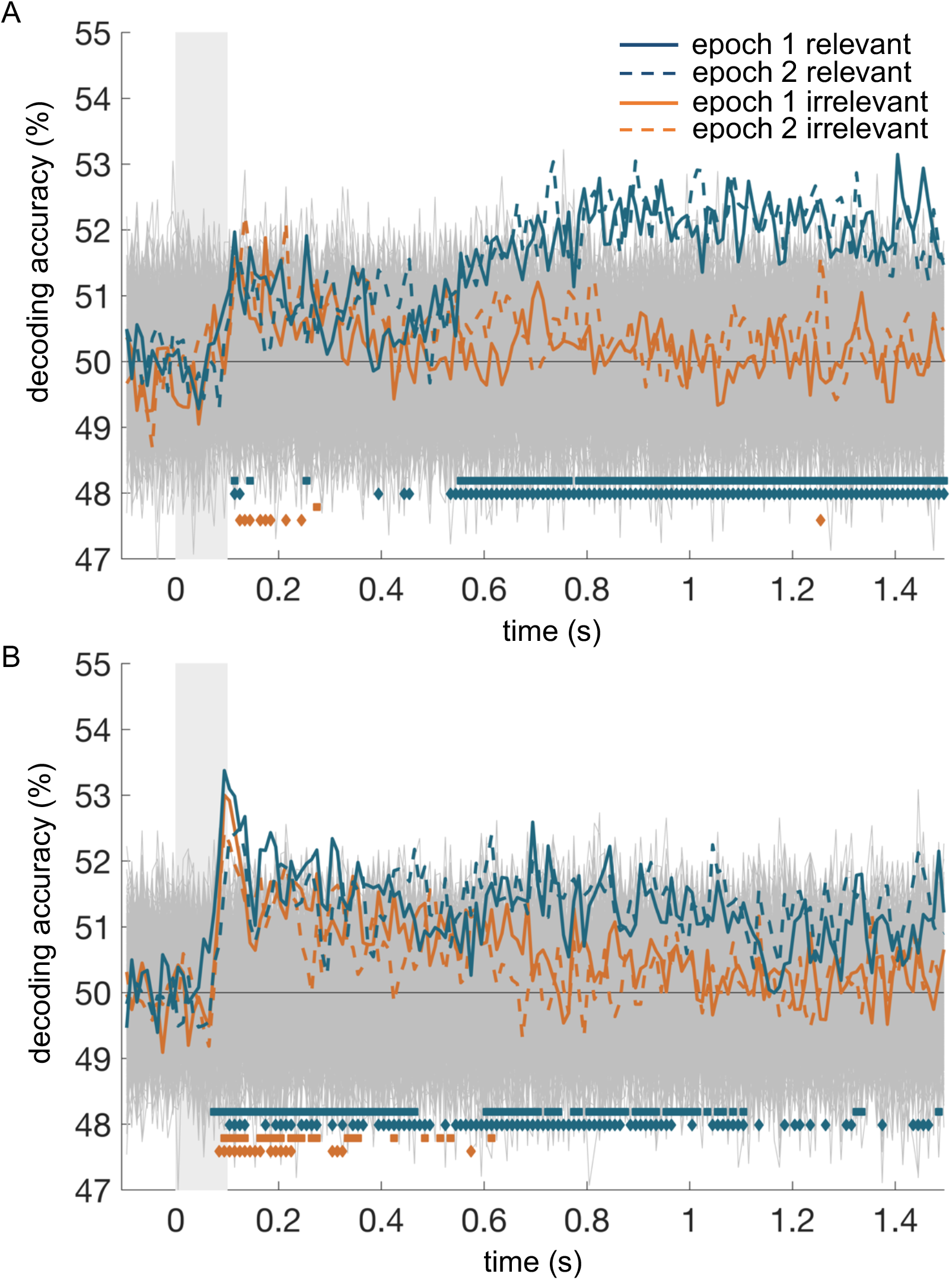
Colour (A) and shape (B) decoding for both epochs superimposed. For each trace, timepoints that reliably differ from chance are marked with coloured squares (solid line, epoch 1) or diamonds (dotted line, epoch 2). There was no reliable difference between epochs, or interaction between epoch and relevance.

## Discussion

Understanding how task-sensitive neural codes reconfigure is a key step in tracing how the brain supports adaptive behaviour. Here, we conducted two experiments to ask whether the brain can rapidly reconfigure neural codes for relevant stimulus features when what is relevant changes. In both experiments, participants judged the shape, then colour, or vice versa, of two targets presented in sequence. When shape and colour judgements were easy (Experiment 1), we observed strong coding of all object information. We found no reliable evidence for preferential coding of task-relevant features. By contrast, when the shape and colour judgements were difficult and additional distractors were present (Experiment 2) we did see preferential coding for the relevant feature. Crucially, stronger coding for the relevant feature occurred in both phases of the trial, even though participants were shifting attention between features mid-trial.

Tracing this process with MEG allows us to see the temporal evolution of preferential coding in the human brain, showing with millisecond resolution how attention emerges and redirects. Even with this precise temporal detail, Experiment 2 demonstrates a remarkably similar timecourse for selection of relevant information for the first and second stimulus. We might expect that preferential encoding of the relevant feature in the second epoch would be slower and/or less selective than in the first. For example, a lag or reduction in selectivity could reflect residual attention to the feature that was relevant for the first epoch, or time taken to transition to selective encoding of the second feature. Instead, we did not find any evidence of slower or reduced selectivity in the second epoch, suggesting that, in this paradigm, reconfiguration was fast enough for the relevant feature of the second stimulus to be selected as efficiently as for the first. These findings indicate that, when adaptive coding is engaged, task-relevant information is preferentially coded with remarkable speed even as task demands change within single trials. This supports the suggestion that goal-directed behaviour involves fast, sub-trial switching of attentional sets (Duncan, 2013).

Although participants successfully performed both tasks, Experiment 1 did not elicit reliably increased neural coding of the relevant stimulus. Curiously, both tasks showed strong and sustained representation of the rule (“attend colour, then shape”), even though only one task showed an effect of rule on stimulus coding. Current explanations of top-down control emphasise both maintaining task information and enhancing relevant stimulus information. For example, both rule and relevant stimulus information can typically be decoded from MD regions in human fMRI (Jackson et al., 2016; Woolgar, Afshar, et al., 2015; Woolgar, Thompson, et al., 2011; Woolgar & Zopf, 2017) and from frontal cortex in non-human primate single-unit recordings (Everling et al., 2006; Stokes et al., 2013). Disrupting prefrontal function causes reduction in task-relevant information coding (Jackson et al., 2021), and incorrect rule or stimulus information coding predicts incorrect behavioural responses (Woolgar et al., 2019). Moreover, the structure of frontal stimulus information predicts subsequent occipital stimulus information as attentional selection of relevant features emerges (Goddard et al., 2019). In view of these findings, it is plausible that selection occurs through rule information that is maintained by domain-general regions, which in turn selectively enhance relevant stimulus information in both domain-general and task-specific regions. In contrast, in Experiment 1 we observed a dissociation: clear rule coding, but no evidence of enhanced coding of the relevant stimulus features, even though the rule defined which stimulus features to attend to. Rule decoding increased after the stimulus displays in both tasks, particularly in Experiment 2. These increases could reflect neural responses diverging as participants applied the feature rule to the stimuli, but if so, this did not appear to be mediated by enhanced processing of the attended stimulus features. Conversely, increases in rule decoding could be related to a more general shift, such as the widespread reduction in cortical response variance at the onset of a stimulus (Churchland et al., 2010). It is important that we do not over-interpret variations in rule decoding. Rather, tracing attentional rule information alongside rule-related changes in stimulus information allows us to more precisely characterise the impact of the rule on attentional selection. Experiment 1 highlights this, showing that more accurate rule decoding after stimulus displays does not necessarily translate to preferential coding of relevant stimulus information.

There were several differences between the two experiments that may have contributed to the different results. Experiment 2 was more difficult: participants responded well above chance level in both tasks, but overall performance was lower in Experiment 2 even after intensive training on the task. In Experiment 1, stimuli were drawn from a set of four objects, with strongly differentiated colours and shapes, and a single object was shown on each display. Because of this small stimulus set, on 25% of trials the objects on Display 1 and Display 2 were identical, making the task trivial. On the remaining trials, participants had to select differential information from each display to respond accurately. However, there was significantly less information on each display, and less confusability among colours and shapes, than in Experiment 2. Thus, responding to the relevant information could well engage different attentional mechanisms across the two tasks.

Increased selection with increased stimulus complexity is a common theme in many theories of attention. For example, behavioural data demonstrate that although participants can find and respond to targets more quickly in simple displays compared to complex displays, they are also more easily influenced by salient distractors (Lavie, 1995; Lavie & Tsal, 1994). Similarly, BOLD activity associated with a distractor stimulus category no longer differentiates repeating and unrepeating distractors when target visibility drops (Yi et al., 2004). Load theory (Lavie, 1995; Lavie et al., 2014), takes these findings to argue that selection is qualitatively different for simple and complex stimuli. In simple environments, perceptual capacity not spent on relevant information “spills over” to other stimuli. As complexity increases, through the number, similarity, or visibility of the stimuli, we voluntarily direct our fixed capacity toward relevant features and ignore salient distractors.

We should note that load theory does not specify how relevant and irrelevant features that fall within perceptual capacity limits are represented to enable accurate responses. For example, we might predict that preferential coding (similar to behavioural responses) only occurs when we exceed our perceptual capacity. Our differential findings in Experiments 1 and 2 could be consistent with this view, provided that Experiment 1 displays fell within most participants’ perceptual capacity while Experiment 2 displays exceeded it. However, neuroimaging data so far do not support the idea that we require complex displays to engage preferential coding. Indeed, multivariate analyses of fMRI data show that relevant feature coding in visual cortex (V1 and LOC) can be enhanced in simple displays, with this enhancement extending to frontoparietal cortex when stimulus discrimination is difficult (Jackson et al., 2016; Woolgar, Williams, et al., 2015). Recent sensor-space MEG data also show enhanced coding of the relevant stimulus category (objects or letters) even though the displays contained only two easily distinguishable objects (Grootswagers et al., 2021). Thus, a better application of load theory to multivariate neuroimaging would predict that feature-selective attention produces a relative enhancement of relevant perceptual information in simple displays, even though both relevant and irrelevant information can be perceived and recalled. This raises an interesting question: if both simple and complex displays can elicit preferential coding (that we can detect with both fMRI and MEG), why is stimulus coding in our Experiment 1 unaffected by relevance?

Theories focusing on the object-based nature of attention (Baldauf & Desimone, 2014; Chen, 2012) may offer a better explanation for why coding two features of a single object, as in our Experiment 1, and coding two objects, as in Grootswagers et al. (2021), would follow different rules. Behavioural studies demonstrate that we can often report irrelevant features of a target object without any apparent performance cost, suggesting that all features of the object are processed in parallel before we chose specific elements to respond to (Chen, 2012; Duncan, 1984). Under this object-based account of attention, it is unsurprising that we did not observe different responses to the same visual feature when it was the relevant or irrelevant dimension of a target object. Rather, we should expect to see preferential coding of the target object over the distractor. We can see this in Goddard et al. (2019), in which a spatial attention effect emerges before coding of the relevant target feature outstrips all other traces. This same pattern is suggested by our secondary analyses, where brief main effects of spatial attention emerge before preferential coding of the relevant target feature (Figure S2, epoch 2 colour and epoch 1 shape). However, object-based accounts struggle to account for preferential coding of single dimensions of stimuli (for example, Jackson et al., 2016; Jackson & Woolgar, 2018), as we observed at later timepoints in Experiment 2.

Biased competition (Desimone & Duncan, 1995; Kastner et al., 1998; J. H. Reynolds et al., 1999) provides a possible unifying framework for the load-driven and object-based characteristics of attention. Similar to load theory, this account proposes that complex stimuli trigger attentional selection. Rather than assuming a threshold for “perceptual capacity”, biased competition suggests that, as distinct representations of stimulus features in early visual cortex feed forward to shared neural populations in higher visual cortex, competition emerges for what feature will be represented at the higher level, forcing selection to occur (Desimone & Duncan, 1995; J. R. Reynolds et al., 2012; Scalf et al., 2013). Because integration co-occurs with broadening receptive fields, even spatially segregated shapes can project to the same neurons and compete for in-depth processing. In our study, the two-object displays of difficult-to-discriminate stimuli in Experiment 2 should elicit more competition than the single-object displays in Experiment 1, creating the opportunity for selection.

Importantly, Duncan (2006) integrates space-, object-, and feature-based attention under the biased competition framework, highlighting that competition drives selection across disparate forms of attention, which can operate independently or in concert. This broader perspective of attention as a family of processes implemented through biased competition has since been embraced by Kravitz and Behrmann (2011), who demonstrate that space-, object-, and feature-based attention can combine to enhance object processing. Combined effects of spatial and feature-based attention have also been observed in nonhuman primates’ lateral intraparietal area (LIP; Ibos and Freedman, 2016). Goddard et al. (2019) similarly show multiplicative effects of spatial and feature-selective attention give rise to selective coding of only the relevant feature at the relevant location. Using the same stimuli, we replicated this finding, showing that coding of the relevant feature at the relevant location is enhanced relative to the irrelevant feature at that location (core analyses; Figure 4) and the distractor features (secondary analyses; see Figure S2).

From a broader perspective, each of these theories touches on an assumption that selection is not always engaged. This “selection-free zone” could be a basic perceptual capacity; object binding that lets us process multiple features of the same object “for free”; or differences in location, colour, and orientation allowing for simultaneous processing without competition. Neural network simulations additionally offer some insight into the cost of selection, showing that strong coding of currently relevant task features induces slow reconfiguration to code subsequently relevant information (Musslick et al., 2018). Therefore, there may be a computational benefit to avoiding re-configuration of attentional sets (e.g., within trials) where possible. An adaptive system may be characterised not only by the ability to flexibly prioritise processing of currently-relevant information, but the flexibility to only do so when processing demands require it.

Here we have shown that human adaptive population codes can reconfigure within a single trial. This supports current theory, which emphasises the potential of focusing on each step in a task to produce complex and creative behaviour. Surprisingly, where attention effects were seen, the dynamics were comparable for between trial and within-trial shifts of attentional focus. This provides a potential neural substrate for the rapid creation of attentional episodes in multi-part tasks. However, significant effects of attention were only obtained in a demanding version of the task. Although many factors differed between the experiments, the difference could reflect the inherent cost of reconfiguring attention, meaning that it is not always an optimal strategy to engage. Future work will be important to identify what conditions push us toward preferentially coding the relevant information. Spatio-temporally resolved methods, such as source reconstructed MEG and MEG-fMRI fusion, paired with systematic manipulation of task difficulty, could further elucidate how domain-general and task-specific brain regions interact to select relevant information under varying task demands. Rapid stimulus streams or self-directed attention shifting could further probe how rapidly the brain can reconfigure neural codes for preferential processing. Furthermore, relating the speed of reconfiguration to measures of fluid ability could clarify the functional importance of adaptive coding timescales. Together with our findings, this will offer rich insight into the biological bases of a mind that adapts to connect our goals with the world around us.

