## supplementaryMaterial for "Neural coding of visual objects rapidly reconfigures to reflect sub-trial shifts in attentional focus"

**Supplementary Material**

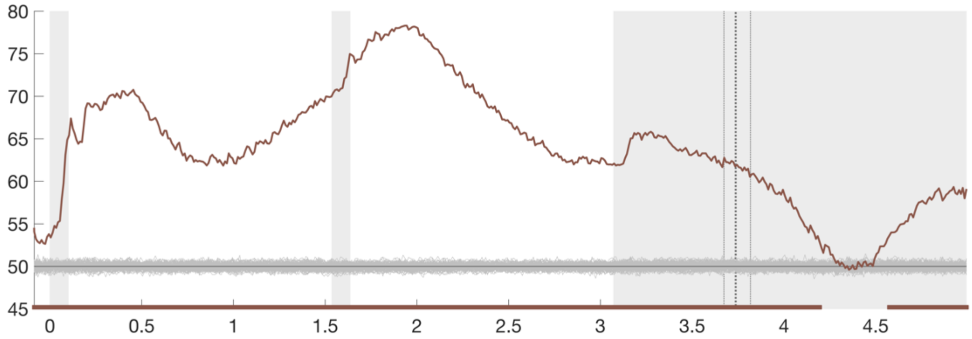


*Figure S1.* Location rule decoding ("attend left then right" vs "attend right then left") for Experiment 2. Vertical grey patches mark the stimulus displays and the maximum possible duration of the choice display. Vertical dotted lines mark the median response time with one quartile on either side. Permutation-based null data are shown in grey around chance (50%). Timepoints at which decoding was reliably different to the null based on threshold-free cluster correction are marked below the trace in brown.

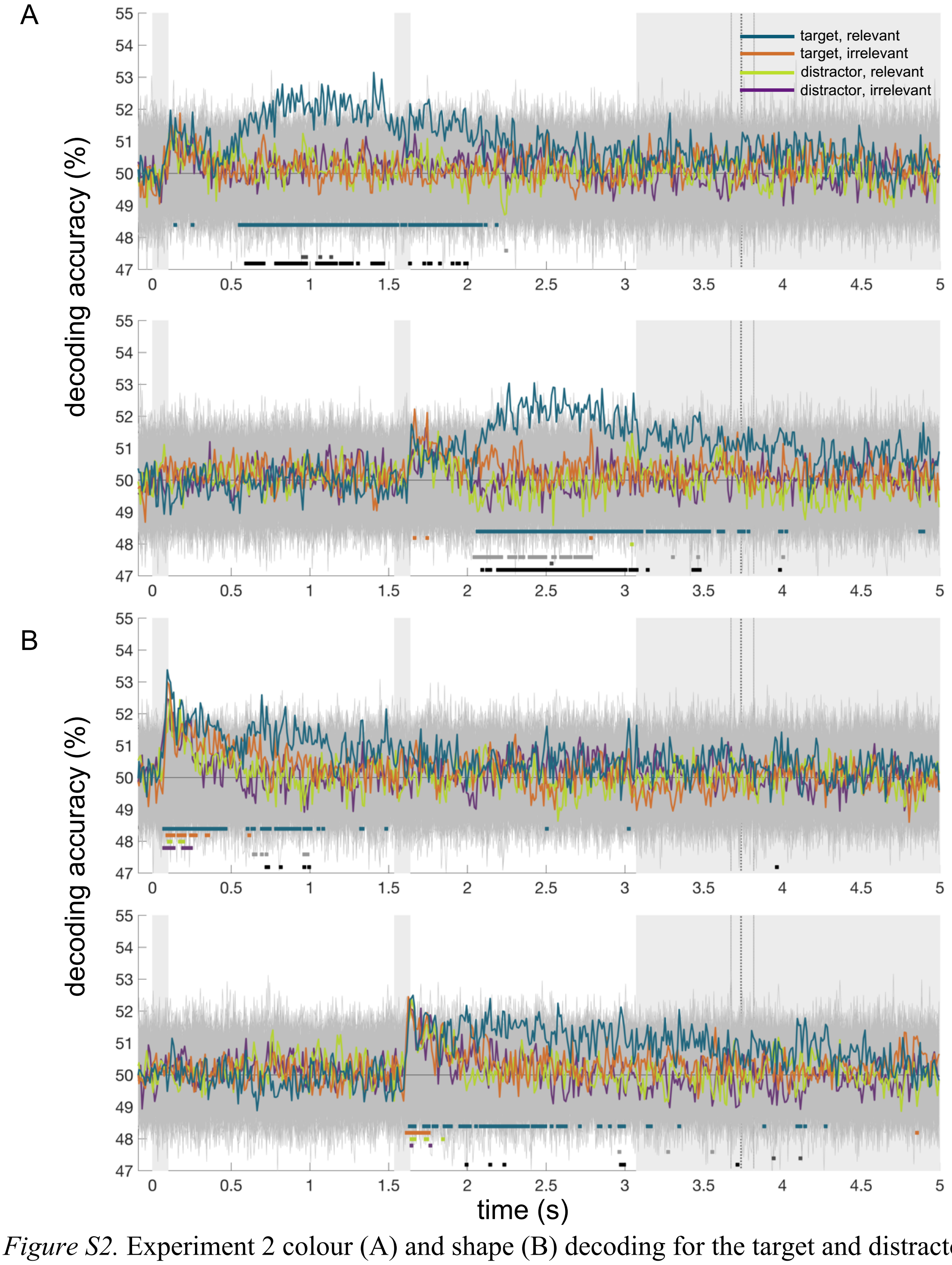


*Figure S2.* Experiment 2 colour (A) and shape (B) decoding for the target and distractor objects on each display. Traces represent decoding accuracy for colours or shapes at the attended location (blue=attended feature, orange=unattended feature) as in Figure 4, as well as at the unattended location (green=attended feature, purple=unattended feature). Times at which each trace was reliably different to chance, at $p<0.05$ with a threshold-free cluster correction for multiple comparisons, are marked in the corresponding colour. Greyscale markers indicate times with a statistically reliable effect of spatial attention (target vs distractor, light grey), feature attention (attended vs unattended feature, dark grey), or interaction between spatial and feature attention (relevant feature of target vs all other features, black).
